# Resting-State EEG Aperiodic Exponent Moderates the Association Between Age and Memory Performance in Older Adults

**DOI:** 10.1101/2025.03.20.643891

**Authors:** Alicia J. Campbell, Toomas Erik Anijärv, Mikael Johansson, Thomas Pace, Jim Lagopoulos, Daniel F. Hermens, Jacob M. Levenstein, Sophie C. Andrews

## Abstract

Memory functions are susceptible to age-related cognitive decline, making it essential to explore the underlying neurophysiological mechanisms that contribute to memory function during healthy ageing. Resting-state EEG (rsEEG) parameters, particularly the aperiodic exponent, a marker of cortical excitation-inhibition balance, and individual alpha peak frequency, a correlate of neural processing efficiency, have demonstrated associations with ageing and cognitive functions. This study investigated associations between these rsEEG markers and performance across multiple memory systems in healthy older adults (n = 99) aged 50-84 years, specifically the direct associations of these markers on memory across episodic, working, and visual short-term memory systems, assessed via computerised tasks, as well as their moderating effects on age-memory relationships. While no direct associations were seen between rsEEG markers and memory performance across tasks beyond the contribution of age, gender and education, results revealed significant moderating effects of the aperiodic exponent on age-related performance in episodic and visual short-term memory. Notably, for individuals with a higher exponent, age was not significantly associated with episodic or visual short-term memory performance, whereas those with average and lower exponent values showed poorer performance with older age. These findings suggest that average and lower aperiodic exponents may reflect a marker of decrement in age-related memory performance and higher exponents may index an underlying protective mechanism against age-related memory decline. This investigation extends the current understanding of cognitive ageing mechanisms by identifying the aperiodic exponent as a potential biomarker explaining individual differences in cognitive ageing trajectories in older adult populations, particularly in episodic and visual short-term memory systems, and establishes a framework for studying neuroprotective mechanisms and developing interventions to preserve cognitive function in older adults.

## 1.0 Introduction

Cognitive decline represents a well-documented feature of normal ageing (Ferguson et al., 2021; Salthouse, 2009), with memory systems exhibiting particularly pronounced vulnerability to age-related deterioration (Nyberg et al., 2012; Raz & Lindenberger, 2011; Salthouse, 2011). Multiple memory systems demonstrate systematic changes across the ageing process: episodic memory, which encompasses the encoding, maintenance, and retrieval of temporally and spatially contextualised personal experiences (Tulving, 2002); working memory, characterised by the active maintenance and manipulation of task-relevant information in the absence of ongoing sensory input (Baddeley, 2012); and visual short-term memory, defined by the capacity-limited temporary storage of visual representations (Logie, 1989; Luck & Vogel, 1997; Phillips, 1974). These memory systems, fundamental to higher-order cognitive functions, demonstrate significant age-associated declines (Brockmole & Logie, 2013; Nyberg et al., 2012; Nyberg & Pudas, 2019; Park et al., 2002; Park & Reuter-Lorenz, 2009; Salthouse, 1991, 2009). Together, these observations underscore the importance of understanding the underlying neurophysiological mechanisms that contribute to memory function throughout the healthy ageing process, particularly among older adults. Such understanding may facilitate identification of factors that promote cognitive resilience or accelerate decline, thereby informing the development of targeted interventions designed to preserve cognitive function in ageing populations.

Electroencephalography (EEG) provides a high-temporal resolution, non-invasive measurement of neural activity that enables detailed investigation of neurophysiological processes. The signals captured through EEG comprise both periodic (oscillatory) and aperiodic (non-oscillatory) components, which together reflect the brain’s overall dynamics. Among the oscillatory activities, alpha rhythms (8–13 Hz) have been extensively studied in the context of cognitive function and ageing-related processes (Babiloni et al., 2006; Başar, 2012; Başar & Güntekin, 2012; Klimesch, 1999; Michels et al., 2013; Scally et al., 2018).

Individual alpha peak frequency (IAF), defined as the peak frequency within an individual’s alpha band, represents a key periodic EEG marker demonstrating associations with ageing and cognitive functions, such as attention (Angelakis et al., 2004; Campbell et al., 2024; Klimesch et al., 2007; Ramsay et al., 2021), memory (Klimesch, 1997; Wianda & Ross, 2019), and information processing speed (Cesnaite et al., 2023; Klimesch et al., 1996). IAF demonstrates systematic developmental modulation across the lifespan, characterised by an increase from childhood to adolescence (Cragg et al., 2011; Klimesch, 1999; Tröndle et al., 2022) and subsequent decline from adulthood into older age (Campbell et al., 2024; Cesnaite et al., 2023; Finley et al., 2022; Ishii et al., 2018; Knyazeva et al., 2018; Mizukami & Katada, 2018; Smith et al., 2023). This modulation with age is thought to reflect underlying neurobiological changes, particularly decreases in white matter integrity associated with ageing (Valdés-Hernández et al., 2010). Individuals with a higher IAF across adulthood and older age demonstrate positive associations with multiple domains of cognitive function, including working memory performance (Cesnaite et al., 2023; R. C. Clark et al., 2004; Klimesch, 1997), sustained attention capacity (Campbell et al., 2024) and cognitive preparedness (Angelakis et al., 2004).

Furthermore, a faster IAF has demonstrated correlations with general intelligence (Grandy, Werkle-Bergner, Chicherio, Lövdén, et al., 2013). While oscillatory activity like IAF provides important insights into cognitive function, non-oscillatory components of neural activity are also crucial to consider.

Regarding the non-oscillatory components of neural activity, the aperiodic activity (sometimes referred to as “background noise”) is characterised by a 1/*f*-like distribution. This distribution reflects a systematic decrease in power as frequency increases. The exponent of this activity corresponds to the negative slope of the power spectrum on a log-log scale and is widely interpreted as reflecting neural excitation-inhibition (E/I) balance (Gao et al., 2017; McKeon et al., 2024; Medel et al., 2023; Podvalny et al., 2015), although this interpretation has also been challenged (Brake et al., 2024; Salvatore et al., 2024). Recent work has shown systematic age-related alterations of the exponent, with older adults generally exhibiting a reduced (flatter) exponent when compared to younger adults (Dave et al., 2018; Merkin et al., 2022; Tran et al., 2020; Tröndle et al., 2023; Voytek et al., 2015) and across the lifespan (Thuwal et al., 2021). However, studies focusing specifically on older adult populations have yielded mixed findings regarding age-associated changes in the exponent. While some studies report a decrease in the exponent within older age ranges (Cesnaite et al., 2023; Finley et al., 2022), others fail to identify significant associations (Campbell et al., 2024; Merkin et al., 2022; Smith et al., 2023). This may be due to sample size, with studies reporting significant age-related effects typically having substantially larger cohorts (Cesnaite et al., 2023; Finley et al., 2022) compared to the studies finding no significant relationships (Campbell et al., 2024; Merkin et al., 2022; Smith et al., 2023), suggesting that age-associated changes in the exponent during later life may represent subtle effects.

Recent research has uncovered connections between the aperiodic exponent and multiple cognitive functions, including lexical prediction (Dave et al., 2018), working memory (Voytek et al., 2015) and interference processing (Kałamała et al., 2024) during task, as well as cognitive speed (Montemurro et al., 2024; Ouyang et al., 2020) and working memory (Thuwal et al., 2021) at rest. However, studies examining resting-state aperiodic activity’s relationship with cognitive function across adulthood and specifically within older populations remain in the early stages and have produced mixed findings.

While several studies have shown associations between the exponent and cognitive performance metrics in older adults, both cross-sectionally (Smith et al., 2023) and longitudinally across the adult lifespan (Finley et al., 2024), other studies have failed to replicate these relationships (Campbell et al., 2024; Cesnaite et al., 2023; Tröndle et al., 2023).

The inconsistency may stem in part from the potential limitations of traditional neuropsychological assessments. Conventional pen-and-paper and verbal assessments are often static, low-dimensional measures of cognition (Pitkow & Angelaki, 2017), such that they typically assess a limited number of constructs with simple stimuli and basic response formats. Furthermore, examiner-administered assessments may introduce potential sources of variability through scoring inconsistencies and examiner bias. In contrast, computerised cognitive assessments offer several enhanced methodological advantages, particularly for investigating cognitive function in ageing populations.

These include increased sensitivity and precision, automated scoring and interpretation, greater standardisation, alternating forms of tasks to minimise practice effects and the ability to measure a broader range of cognitive processes (Gates & Kochan, 2015; Parsons et al., 2018; Wild et al., 2008; Zygouris & Tsolaki, 2015).

In our previous study, which examined resting-state EEG (rsEEG) correlates of sustained attention in healthy older adults (50–84 years), we identified distinct relationships between neurophysiological parameters and performance on computerised sustained attention tasks (Campbell et al., 2024). IAF emerged as a key neural correlate, exhibiting positive associations with performance on the Sustained Attention to Reaction Task (SART; Robertson et al., 1997) and mediated age-related performance effects. However, our previous investigation did not examine memory function within the studied cohort. The current literature investigating relationships between rsEEG IAF, the exponent and memory performance within healthy older adult cohorts is limited, demonstrates conflicting findings and is primarily based on traditional non-computerised neuropsychological assessments (Cesnaite et al., 2023; Finley et al., 2024; Smith et al., 2023; Tröndle et al., 2023). This represents a significant gap in understanding the relationship between rsEEG markers and memory function in older age.

To address this gap and extend our previous work on rsEEG correlates of sustained attention in a healthy older cohort, the present study conducted an exhaustive investigation on how the aperiodic exponent, IAF and age relate to performance on a battery of computerised memory tasks using the Cambridge Automated Neuropsychological Test Battery (CANTAB; Cambridge Cognition Ltd., 2015), targeting different memory systems in the same sample of older adults. More specifically, the present study aimed to: (i) explicate direct associations between the rsEEG measures and performance across multiple computerised memory systems in healthy older adults; (ii) assess potential moderating effects of rsEEG measures on age-memory performance relationships.

We hypothesised positive associations between memory performance across all tasks and both IAF and the 1/*f* exponent. Furthermore, we anticipated significant interaction effects between age and these rsEEG markers in predicting memory performance across all tasks. Specifically, we hypothesised that the relationship between age and performance on each memory task would be more pronounced (e.g., poorer memory performance with advancing age) at lower exponent and IAF values. By incorporating both oscillatory and aperiodic rsEEG markers in conjunction with sensitive, computerised memory assessments, this study sought to advance the current understanding of the relationships between neural dynamics, age and memory function in healthy older adults.

## 2.0 Methods

### 2.1 Participants

The cross-sectional analysis in this study utilised data from 99 healthy individuals aged 50 to 84 years from the Lifestyle Intervention Study for Dementia Risk Reduction (LEISURE). LEISURE, an ongoing study based at the University of the Sunshine Coast’s Thompson Institute, has its full protocol published separately (Treacy et al., 2023). In summary, participants were recruited from the community, aged 50-85, fluent in English, and generally in good health. Exclusion criteria included diagnosed psychiatric or major neurological conditions, mild cognitive impairment, significant head trauma history (loss of consciousness>60 min), stroke, epilepsy, or current use of psychotropic drugs. The study’s protocol received approval from the University’s Human Research Ethics Committee (A191301) and was registered with the Australian New Zealand Clinical Trials Registry (ACTRN12620000054910). All participants provided written informed consent before their involvement in the study.

### 2.2 Experimental Procedure

As previously published (Campbell et al., 2024), participants first completed the CANTAB battery. The rsEEG assessment was conducted at the start of the EEG session, which generally occurred the next day or, if not possible, within the same week following the CANTAB battery.

### 2.3 Cognitive Battery

Participants were individually assessed using the CANTAB (Cambridge Cognition Ltd., 2015) on an iPad touchscreen interface. Three neuropsychological tasks were selected from the battery completed by the participants to evaluate memory functions: Delayed Matching to Sample (DMS), Paired Associates Learning (PAL) and Spatial Working Memory (SWM).

The DMS task, lasting approximately 7 minutes, assesses visual simultaneous and short-term memory. Participants are presented with a complex visual pattern and then asked to select the matching pattern from four similar options. These options are displayed either simultaneously or after a delay of 0, 4, or 12 seconds following the disappearance of the original pattern. The primary outcome measure is Percent Correct All Delays (DMS PCAD), representing the percentage of assessment trials containing a delay during which the subject chose the correct box on their first box choice. Higher scores indicate better performance.

The PAL task, lasting approximately 8 minutes, assesses visuospatial episodic memory and learning. Participants are presented with containers arranged around the edge of the display, which open randomly to reveal complex visual patterns. Subsequently, the patterns appear one by one in the centre of the screen, and participants must select the original container associated with each pattern. The difficulty level increases progressively from 2 to 8 containers. Participants must complete each level without errors to progress; failing four times at any level ends the test. The primary outcome measure is Total Errors Adjusted (PAL TEA), which is the total number of errors across all assessed trials and stages, adjusted for incomplete or failed trials (assuming 95% error and 5% correct by chance for remaining stages). Lower scores indicate better performance.

The SWM task, lasting approximately 4-6 minutes, assesses spatial working memory. Participants are presented with a series of boxes and must tap them to find a hidden token. In each trial, participants must avoid revisiting boxes where a token has already been found. The number of boxes increases progressively from four to six and finally to eight. The colour and position of the boxes vary with each trial to prevent learning effects. The primary outcome measure is Total Between-Search Errors (SWM BE), which is the number of times the participant returns to a box in which a token has already been found across all trials. Lower scores indicate better performance.

### 2.4 Electrophysiological Recordings

rsEEG data were acquired using a 32-channel Active-Two BioSemi system (BioSemi Active-Two, V.O.F., Amsterdam, Netherlands) during four minutes of eyes-closed resting state. For comprehensive details of the data acquisition protocol, readers are referred to Campbell et al. (2024).

### 2.5 Electroencephalographic Analysis

#### 2.5.1 Signal Pre-processing

Signal preprocessing followed procedures are detailed in Campbell et al., (2024). Briefly, the data underwent offline referencing to the common average reference, band-pass filtering (1-30 Hz), artifact rejection, using MNE Python (Gramfort et al., 2013), an open-source software package for EEG/MEG analysis and Autoreject (Jas et al., 2016, 2017), a tool for automated artifact rejection, as well as segmentation into 5-second epochs without overlap.

#### 2.5.2 Spectral Analysis

The spectral analysis involved two main steps: calculating the power spectrum density (PSD) of the rsEEG signals and decomposing the PSD into periodic and aperiodic signal components. Initially, the preprocessed rsEEG data were converted to the frequency domain by estimating PSDs using Welch’s method (Welch, 1967). This was done for the 1-30 Hz range, employing a 2.5-second Hamming window with 50% overlap and 97.5-second zero-padding. The extended zero-padding allowed for finer interpolation in the frequency domain, enabling a more precise estimation of IAF to 0.1 Hz accuracy while acknowledging this doesn’t increase true frequency resolution. Average PSD values (µV2/Hz) were calculated in bilateral parieto-occipital (P3, Pz, P4, PO3, PO4, O1, Oz, O2) and bilateral fronto-central (F1, Fz, F2) electrode clusters. The parieto-occipital electrode cluster was selected based on prior scientific literature, which highlights prominent age-related patterns in the alpha domain (Babiloni et al., 2006; Grandy, Werkle-Bergner, Chicherio, Schmiedek, et al., 2013; Klimesch, 1999; Knyazeva et al., 2018; Mizukami & Katada, 2018; Rossini et al., 2007; Scally et al., 2018; Sghirripa et al., 2021) and aligning with recent studies employing similar measures (Donoghue et al., 2020; Montemurro et al., 2024; Tröndle et al., 2022, 2023). The fronto-central electrode cluster selection was guided by emerging evidence demonstrating its significance in cognitive ageing processes (Khalilian et al., 2024) and aligns with recent studies employing similar spectral measures (Cesnaite et al., 2023; Finley et al., 2024; Montemurro et al., 2024). Additionally, a recent study has established an association between aperiodic activity in frontal regions and declining performance in processing speed and executive function domains (Pathania et al., 2022).

Subsequently, the ‘specparam’ toolbox (Donoghue et al., 2020) was used to isolate the aperiodic component of the spectra, specifically the aperiodic exponent, from the log-transformed regional PSDs. The ‘fixed’ aperiodic mode was selected based on visual inspection of the log-log space output, which showed no distinct ‘knee’ in the power spectrum. The algorithm settings were: peak width limits (1-12 Hz), unlimited maximum number of peaks, minimum peak height (0.05 dB), peak threshold (2 standard deviations), and fixed aperiodic mode. Parameterisation was performed across the 1-30 Hz frequency range. While the exponent was the focus of the analyses, the aperiodic offset (y-intercept of the EEG broadband), was also extracted. The final step involved flattening the spectrum by removing the aperiodic component from the original spectrum. Within the flattened spectrum, the peak with maximum power was identified in the 7-14 Hz range, and the IAF was extracted.

Preprocessing and spectral analysis were implemented using a custom-developed EEG data processing pipeline. The specific code is publicly available in the EEG-notebooks GitHub repository (https://github.com/aliciajcampbell/EEG-notebooks).

#### 2.5.3 Exclusion Criteria

Prior to statistical analyses, rsEEG data were excluded based on the following criteria:

*1. Insufficient artifact-free data.*

If the signal contained less than 50% artifact-free data after global and/or local artifact rejection, i.e., < 24 remaining epochs out of the total 48 (n = 1).

*2. Poor spectral fit.*

If the fit of the parameterised power spectrum to the original PSD was below R² < 0.9 (n=3 in the fronto-central electrode cluster) or contained significant omissions of signal, as determined by visual inspection (n=4 in the fronto-central electrode cluster and n=3 in the parieto-occipital cluster).

*3. Absence of alpha peak.*

If no alpha peak was detected, the IAF measure was excluded from the analyses. However, the aperiodic component was retained (n=3 with no peaks in the fronto-central electrode cluster and n=3 with no peaks in the parieto-occipital cluster).

### 2.6 Statistical Analyses

Statistical analyses were performed using Python scripts, utilising: Pandas (McKinney, 2010; The pandas development team, 2023), NumPy (Harris et al., 2020), SciPy (Virtanen et al., 2020), Pingouin (Vallat, 2018), Statsmodels (Seabold & Perktold, 2010), HLR (Anijärv et al., 2024), Matplotlib (Caswell et al., 2023; Hunter, 2007), Seaborn (Waskom, 2021) and Statannotations (Charlier et al., 2022).

Outliers were identified using z-scores, such that z-scores >3 or <3 were considered extreme cases. Outlier adjustment was performed as required using the z-score standard deviation transformation method, whereby extreme cases were adjusted to one unit above or below the nearest value existing within acceptable z-score ranges (Tabachnick & Fidell, 2013). One univariate outlier was adjusted in the IAF data in the fronto-central electrode cluster, one univariate outlier was adjusted in the spatial working memory errors (SWM BE) data and one univariate outlier was adjusted in the education data. Skewness and kurtosis values for raw and outlier-corrected variables are provided in Supplementary Materials.

Gender and handedness were coded as binary variables, with gender coded as 0 for male and 1 for female, and handedness coded as 0 for non-right-handed and 1 for right-handed. Continuous variables were z-scored for statistical analysis.

Descriptive statistics for continuous variables were reported as mean, standard deviation, and range (minimum-maximum), and categorical variables were reported as N. To assess the pattern of associations among variables of interest, bivariate correlations were calculated between age, education, memory task scores (DMS PCAD, PAL TEA, SWM BE) and rsEEG measures (IAF and aperiodic exponent at fronto-central and parieto-occipital electrodes) using Spearman’s rank (two-tailed). A total of 36 pairwise comparisons were conducted. To control for multiple comparisons, *p*-values of the 36 comparisons were corrected using the false discovery rate (FDR) method at q = .05 (Benjamini & Hochberg, 1995). Statistical significance was determined using an alpha level of .05.

The association between rsEEG measures and performance across each memory task was examined using independent hierarchical linear regression (HLR) models. Each model evaluated the direct association between a specific rsEEG measure, either the aperiodic exponent or IAF in either the fronto-central or parieto-occipital electrode cluster, and performance on memory tasks. Next, to assess the potential moderation of age-related differences in memory performance by rsEEG measures, interaction terms between age and the relevant rsEEG measure were also examined. Each HLR model used a memory task score (either DMS PCAD, PAL TEA, SWM BE) as the dependent variable and included a single rsEEG predictor. In constructing the HLR models, demographic covariates, including age (at the time of EEG recording), gender, years of education and handedness were controlled for in the first step. The rsEEG measure of interest was added in the second step, and the interaction term (Age × rsEEG measure) was introduced in the third step. Improvements in model fit were assessed using the F-change statistic (ΔF) and its associated p-value. Statistical significance was determined using an alpha level of .05.

Given findings that show the aperiodic offset declines with age (Cesnaite et al., 2023; M. Clark et al., 2024; Finley et al., 2024; Merkin et al., 2022; Voytek et al., 2015) and similar to the exponent, has been found to be predictive of cognitive performance (Euler et al., 2024; Immink et al., 2021; McKeown et al., 2024; Ouyang et al., 2020), additional analyses of the aperiodic offset were conducted using the same procedures as described above. Detailed results are provided in the Supplementary Materials.

To further explore significant interaction terms identified in the HLR models, simple slope analyses were conducted. These analyses examined the relationship between age and task performance at three levels of the individual rsEEG measure: low (≤−1 SD below the mean), mean (±1 SD) and high (≥+1 SD above the mean). The simple slopes were calculated using the regression coefficients for the z-scored variables and the significant interaction term, with standard errors derived from the covariance matrix.

## 3.0 Results

### 3.1 Descriptive Analyses

We have analysed data of 99 participants (M*_age_* = 65.47, SD = 8.3; 80 females). Demographic and sample information can be seen in Table 1. A histogram illustrating the age distribution of participants is provided in Supplementary Materials.

**Table 1.** Sample descriptives.

| Variable | N | M (SD) | Min, Max |
| --- | --- | --- | --- |
| Age (years) | 99 | 65.47 (8.30) | 50.16, 84.04 |
| Gender |  |  |  |
| Female | 80 |  |  |
| Male | 19 |  |  |
| Education (years) | 99 | 14.78 (2.84) | 9.00, 19.00 |
| Handedness |  |  |  |
| Right | 86 |  |  |
| Left/ambidextrous | 13 |  |  |
| DMS PCAD | 99 | 83.44 (12.02) | 47.00, 100.00 |
| PAL TEA | 99 | 19.60 (14.63) | 0.00, 57.00 |
| SWM BE | 99 | 17.40 (7.47) | 0.00, 30.00 |
| *Fronto-central electrode cluster* |  |  |  |
| IAF | 88 | 9.64 (1.15) | 7.00, 12.32 |
| Exponent | 91 | 0.94 (0.21) | 0.42, 1.56 |
| *Parieto-occipital electrode cluster* |  |  |  |
| IAF | 92 | 9.74 (1.09) | 7.03, 11.9 |
| Exponent | 95 | 0.93 (0.19) | 0.50, 1.36 |

M, mean; SD, standard deviation; DMS PCAD, Delayed Match to Sample percent correct (all delays, visual short-term memory); PAL TEA, Paired Associates Learning total errors (adjusted, visuospatial episodic memory); SWM BE, Spatial Working Memory between errors; IAF, Individual alpha peak frequency.

Bivariate correlations, shown in Figure 1, revealed significant associations between age and performance measures across the memory tasks. Specifically, age exhibited significant positive associations with visuospatial episodic memory errors (PAL total errors, adjusted) and spatial working memory errors (SWM between errors), suggesting poorer performance across these tasks with increasing age. Age exhibited a negative association with visual short-term memory accuracy (DMS percent correct, all delays), however, this association did not survive correction for multiple comparisons. Among the scores across memory tasks, spatial working memory errors showed a positive association with visuospatial episodic memory errors. No other significant associations were observed between the memory task scores.

**Figure 1.**
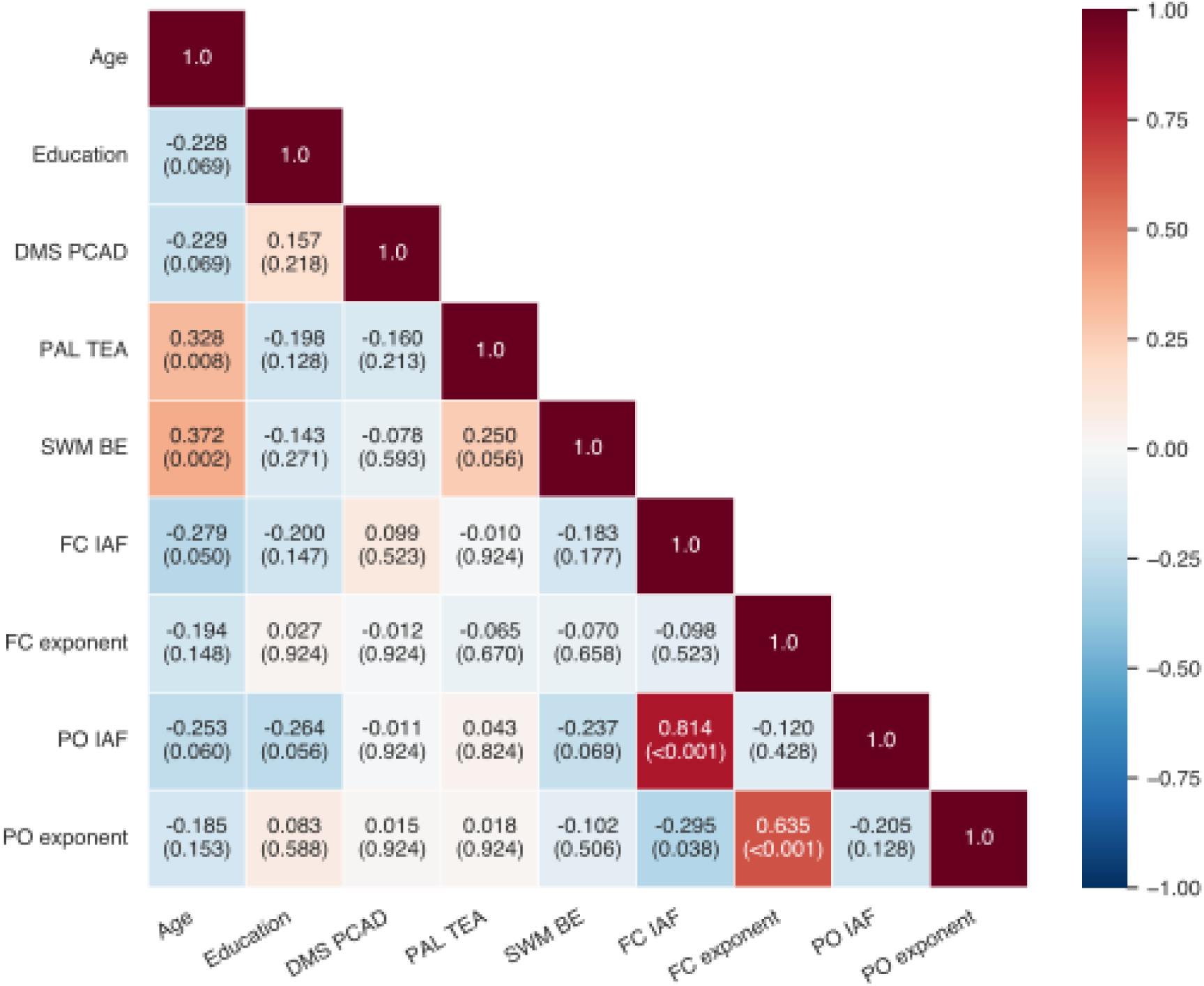
Spearman rho correlation coefficients (ρ) (coloured by magnitude) between performance across memory tasks, age, education and resting-state rsEEG measures, with p-values provided in parentheses; p-values were corrected using false discovery rate (FDR) at .05 across all pairwise comparisons. Statistical significance was determined using an alpha level of .05; DMS PCAD, Delayed Match to Sample percent correct (all delays, visual short-term memory); PAL TEA, Paired Associates Learning total errors (adjusted, visuospatial episodic memory); SWM BE, Spatial Working Memory between errors; FC, fronto-central; PO, parieto-occipital; IAF, Individual alpha peak frequency.

Regarding the rsEEG measures, IAF in the fronto-central electrode cluster showed a significant negative association with age. Similarly, IAF in the parieto-occipital cluster showed a negative association with age, however, this association did not survive correction for multiple comparisons. The exponent in both electrode clusters did not exhibit significant age-related associations. Additional scatter plots visualising the association between age and rsEEG markers are provided in Supplementary Materials. Within the rsEEG measures, IAF in the fronto-central cluster exhibited a negative association with the exponent in the parieto-occipital cluster. Additionally, IAF values demonstrated significant positive correlations between fronto-central and parieto-occipital electrode clusters. Similarly, the exponent showed significant positive correlations between fronto-central and parieto-occipital electrode clusters.

Regarding rsEEG and memory performance relationships across tasks, initially, IAF in the parieto-occipital electrode cluster showed an association with spatial working memory error scores, however, this association also did not survive correction for multiple comparisons. Thus, no significant associations were observed between rsEEG measures and memory performance across tasks.

Figure 1. displays bivariate correlations with p-values corrected for multiple comparisons. For reference, a figure with the uncorrected p-values are provided in Supplementary Materials.

### 3.2 Hierarchical Linear Regressions

#### 3.2.1 Direct Associations

To examine if the rsEEG measures of interest were directly associated with performance across memory tasks after controlling for demographic variables entered in step 1, we analysed the rsEEG measure added in step 2 of the HLR models.

For fronto-central rsEEG measures, the addition of IAF in step 2 did not significantly improve the overall model fit for visual short-term memory accuracy scores (DMS percent correct, all delays, ΔF = .097, *p* = .756), visuospatial episodic memory error scores (PAL total errors, adjusted, ΔF = .026, *p* = .872) or spatial working memory error scores (SWM between errors, ΔF = .306, *p* = .581). Similarly, the addition of the exponent in step 2 did not significantly improve the overall model fit for visual short-term memory accuracy scores (ΔF = .881, *p* = .350), visuospatial episodic memory error scores (ΔF = .107, *p* = .745) or spatial working memory error scores (ΔF = .063, *p* = .802).

For parieto-occipital rsEEG measures, the addition of IAF at step 2 showed no significant predicted improvement for visual short-term memory accuracy scores (ΔF = .096, *p* = .758), visuospatial episodic memory error scores (ΔF = 1.032, *p* = .313) or spatial working memory error scores (ΔF = 1.825, *p* = .180). Similarly, the exponent did not significantly enhance model prediction for visual short-term memory accuracy scores (ΔF = .301, *p* = .584), visuospatial episodic memory error scores (ΔF <.001, *p* = .997) or spatial working memory error scores (ΔF = .009, *p* = .924).

Collectively, neither rsEEG measure in each electrode cluster contributed significantly to explaining memory performance in each task after adjusting for demographic variables in the models.

#### 3.2.2. Interaction Effects (Moderation)

To examine whether the rsEEG measures of interest moderated age-related differences in performance across memory tasks, HLR models assessed moderation effects by testing age × rsEEG interactions added in step 3.

For fronto-central measures, the age × exponent interaction term showed a significant positive association with visual short-term memory accuracy scores (DMS percent correct all delays, standardised β = .221, *p* = .034), accounting for a 4.7% increase in explained variance and significantly improved the overall model fit (ΔF = 4.626, *p* = .034). This suggests that the fronto-central exponent moderates age effects on visual short-term memory accuracy. Notably, no significant moderation emerged for fronto-central exponent effects on visuospatial episodic memory error (PAL total errors, adjusted, ΔF = .723, *p* = .398) or spatial working memory error (SWM between errors, ΔF = .259, *p* = .612) scores, nor for fronto-central IAF interactions across visual short-term memory accuracy (ΔF = .591, *p* = .444), visuospatial episodic memory error (ΔF = .665, *p* = .417) or spatial working memory error (ΔF = .222, *p* = .639) scores.

For parieto-occipital measures, the age × exponent interaction showed a significant negative association with visuospatial episodic memory error scores (standardised β = −.220, *p* = .028), accounting for a 4.6% increase in explained variance and significantly improved the overall model fit (ΔF = 4.997, *p* = .028). This suggests that the parieto-occipital exponent moderates age effects on visuospatial episodic memory errors. Notably, no significant moderation occurred for parieto-occipital exponent effects on visual short-term memory accuracy (ΔF = 3.261, *p* = .074) and spatial working memory error scores (ΔF = .054, *p* = .818). Parieto-occipital IAF interactions failed to reach significance for visual short-term memory accuracy (ΔF = .251; *p* = .617), visuospatial episodic memory error (ΔF = 1.237, *p* = .269) or spatial working memory error (ΔF = 2.671, *p* = .106) scores.

Significant models can be seen in Table 2, while statistics for all models are provided in Supplementary Materials for further reference.

**Table 2.**
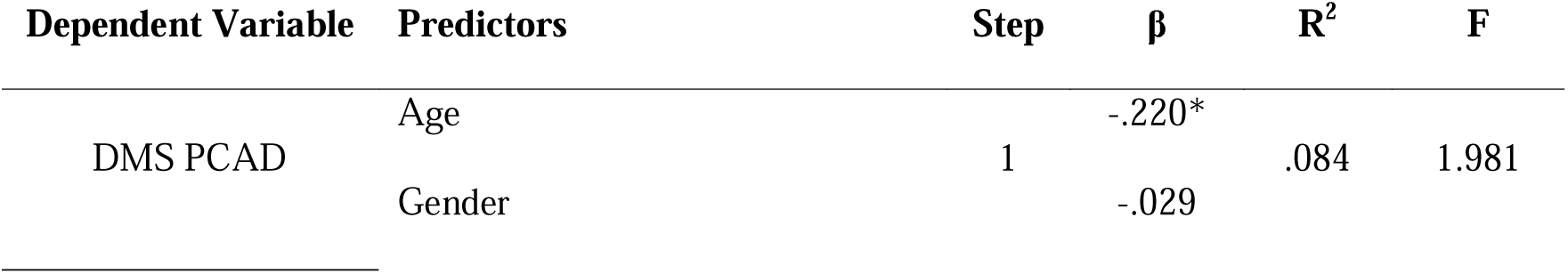
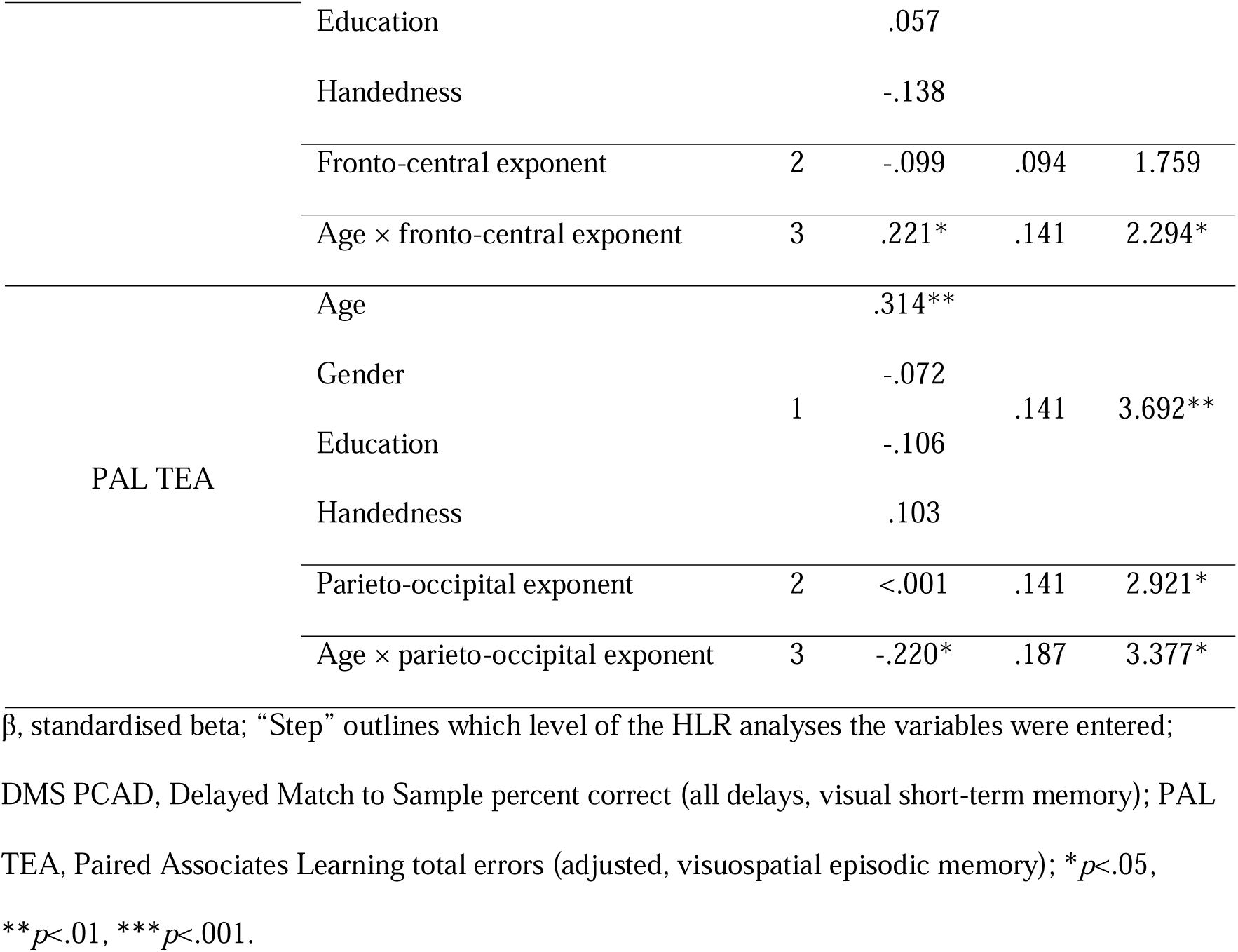
Hierarchical linear regression models predicting memory task performance from main effects of fronto-central IAF and aperiodic exponent values and interactions with age, including demographic covariates as step 1 regressors (significant models only).

### 3.3 Simple Slope Analyses

Simple slope analyses probed the conditional effects of age on memory tasks performance at distinct levels (low, ≤−1 SD below the mean; mean, ±1 SD; high, ≥+1 SD above the mean) of the moderating rsEEG measures identified in step 3 of the significant HLR models.

For the fronto-central exponent’s moderation of visual short-term memory accuracy scores (DMS percent correct, all delays), age showed significant negative associations at low (*p =* .003) and mean (*p =* .019) exponent levels, but not at high levels (*p =* .877), suggesting that the negative effect of age on visual short-term memory accuracy attenuates as the fronto-central exponent increases (Table 3; Figure 2A). Similarly, for the parieto-occipital exponent’s moderation of visuospatial episodic memory error scores (PAL total errors, adjusted), age showed significant positive associations at low (*p =* <.001) and mean (*p =* .002) exponent levels, with no significant age effects observed at high levels (*p* = .535), suggesting that the positive effect of age on visuospatial episodic memory errors attenuates as the parieto-occipital exponent increases (Table 3; Figure 2B).

**Figure 2.**
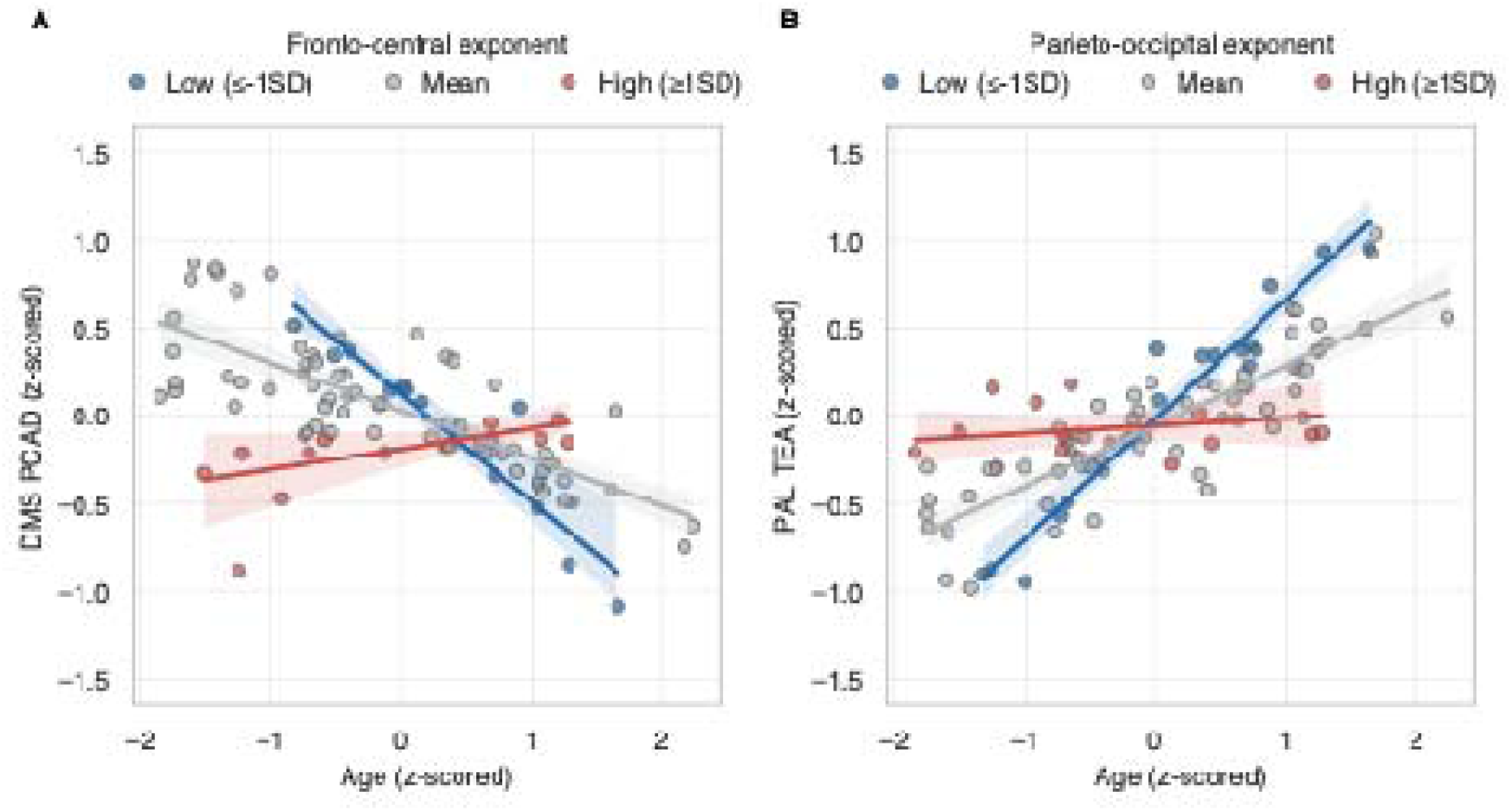
Age associations with memory task performance moderated by resting-state aperiodic exponents. (A-B) Scatter plots with regression lines show relationships between age (z-scored, x-axis) and model-predicted task performance (y-axis; hierarchical linear regression, step 3). (A) DMS PCAD (visual short-term memory) predicted scores. (B) PAL TEA (visuospatial episodic memory). Coloured lines indicate simple slopes for aperiodic exponent groups: Low (≤−1SD, blue), Mean (±1 SD, grey), and High (≥1SD, red) in fronto-central (A) and parieto-occipital (B) electrode clusters. Shaded bands = 95% confidence intervals. DMS PCAD, Delayed Match to Sample percent correct (all delays); PAL TEA, Paired Associates Learning total errors (adjusted).

**Table 3.**
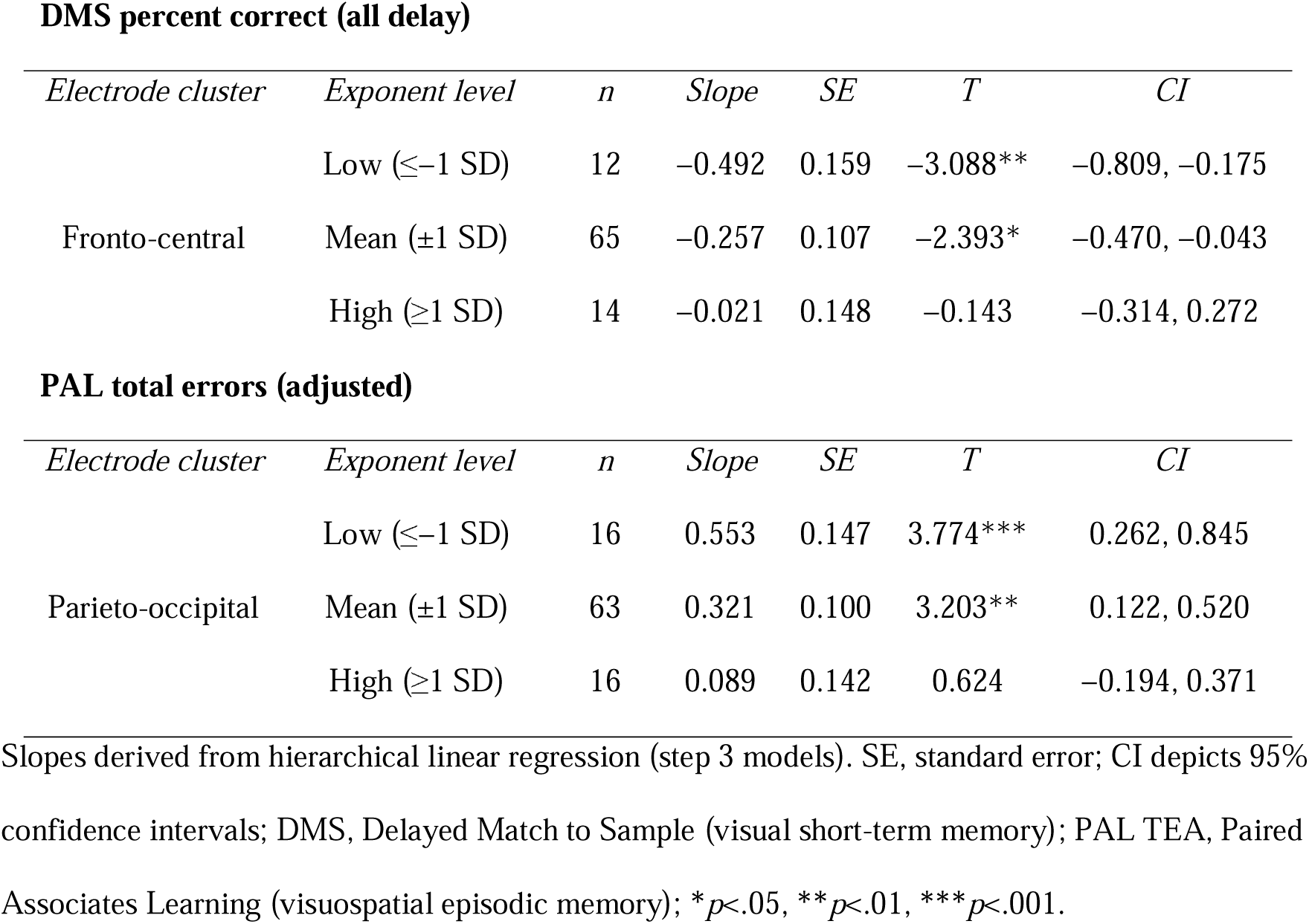
Simple slopes analysis results for age-related associations in memory task performance moderated by low (≤−1SD), mean (±1 SD), and high (≥1 SD) levels of the exponent.

Together, these patterns highlight the moderating role of the fronto-central exponent in the relationship between age and visual short-term memory performance, as well as the moderating role of the parieto-occipital exponent in the relationship between age and visuospatial episodic memory performance.

## 4.0 Discussion

The present study aimed to investigate how the aperiodic exponent and IAF, relate to memory performance in healthy older adults. Specifically, we examined whether these neural measures were associated with performance on computerised tasks from the CANTAB battery that target various memory systems, including visual short-term memory (DMS), episodic memory (PAL) and spatial working memory (SWM), and whether they moderated the relationship between age and memory performance. While we found no evidence for direct associations within the HLR models, moderation analysis via interaction terms in the HLR’s and simple slope analyses revealed that the aperiodic exponent moderated the association between age and performance on the visual short-term memory and visuo-spatial episodic memory tasks. Specifically, within this cross-sectional sample, older adults with a lower exponent showed a stronger negative association between age and memory performance, whereas those with a higher exponent showed a less pronounced age-related association. Notably, the associations between the aperiodic exponent and memory performance differed depending on the electrode cluster: the fronto-central aperiodic exponent was selectively related to visual short-term memory performance, whereas the parieto-occipital aperiodic exponent was related to episodic memory performance.

Research largely indicates that neural network dynamics, specifically E/I balance, governed by glutamate-mediated excitation and GABA-mediated inhibition, can be indexed by the aperiodic exponent of the PSD (Gao et al., 2017; Hong & Rebec, 2012; Waschke et al., 2017). The power distribution in local field potentials (LFPs) is suggested to reflect this dynamic, such that a steeper or increased exponent represents inhibitory activity dominating over excitatory, and a flatter or decreased exponent represents a shift toward excitation over inhibition (Donoghue et al., 2020; Gao et al., 2017; Wang, 2020). However, challenging this interpretation, recent modelling suggests that the exponent may also be shaped by other biophysical factors, such as synaptic time scales and leakage currents (Brake et al., 2024), and a pharmacological study manipulating E/I balance in mice found that while increased inhibition reliably elevates the exponent, increased excitation does not consistently reduce it (Salvatore et al., 2024). Conversely, a recent magnetic resonance spectroscopy study demonstrated that steeper exponents were significantly associated with greater glutamate-GABA asymmetry, primarily driven by glutamate levels (McKeon et al., 2024). This association validates the exponent’s utility as an index of E/I dynamics, with particular sensitivity to excitatory neural processes. Therefore, while some mechanistic ambiguity persists, the convergence of current evidence nonetheless positions the aperiodic exponent as a meaningful, if nuanced, indicator of neural E/I dynamics. Interpreting our results through this framework, we suggest that the rsEEG exponent represents a potential biomarker for individual variability in age-related memory trajectories, as it likely reflects underlying shifts in E/I balance that modulate cognitive ageing outcomes. Specifically, we suggest that lower exponents, potentially reflecting an excitation-dominant neural state, may signal increased vulnerability to age-related memory decline, particularly in memory systems such as episodic and visual short-term memory where declines are well-documented (Adamowicz, 1976; Brockmole & Logie, 2013; Menegaux et al., 2020; Mitchell & Cusack, 2018; Tromp et al., 2015). By contrast, higher exponents, potentially indicative of more robust inhibitory regulation, may index an underlying protective mechanism against these particular age-associated declines.

It is thought that neural homeostasis in cortical networks requires precise regulation of E/I synaptic transmission (Roudi & Latham, 2007; Shew et al., 2011), serving as a crucial mechanism that governs information processing and behavioural output within the central nervous system (Vogels et al., 2011). Disruptions to this balance, characterised by an increase in glutamatergic transmission or diminished GABAergic inhibition, may precipitate brain circuit and large-scale network dysfunction such as heightened neuronal network excitability, potentially compromising social functioning and cognitive performance across multiple domains (López-Sanz et al., 2017; Mendez et al., 2018; Stam et al., 2009). A growing body of research suggests that maintenance of optimal E/I dynamics promotes efficient information encoding (Denève & Machens, 2016), transmission (Shew et al., 2011) and information processing (Kinouchi & Copelli, 2006; Markram et al., 2004; Yizhar et al., 2011), processes fundamental to both episodic and visual short-term memory. The present study’s finding that the aperiodic exponent moderates the association between age and memory performance in older adults highlights its potential role as a neurophysiological marker explaining individual differences in cognitive ageing trajectories, particularly in episodic and visual short-term memory systems, and offers a potential target for interventions aimed at maintaining memory function in later life. Given the cross-sectional design of the current study, future longitudinal research is warranted to track exponent trajectories in relation to age-related memory decline and investigate causality.

Interestingly, recent studies have reported conflicting findings that differ from ours regarding the interplay between the aperiodic exponent, age, and memory. A large-scale investigation (n = 1073) by Cesnaite et al., (2023) examining older adults (60-80 years) in a similar age range to the present study (50-84 years) failed to demonstrate significant modulatory effects of the exponent on episodic memory performance, both directly and through interaction with age. The episodic memory measure in this study was derived from an exploratory factor analysis of various cognitive scales. Similarly, Finley et al., (2024) explored the aperiodic exponent and a factor representing episodic memory at baseline or over a ten-year period in participants aged 36–83 (n = 235) and while the interaction between age and exponent on cognitive performance was not tested, reported no association between the aperiodic exponent and episodic memory, though they did find a relationship with executive functioning. A notable methodological difference emerges in the assessment approaches, such that these prior studies employed traditional neuropsychological assessments, such as pen-and-paper and verbal tests, and performance in specific domains quantified through factorial analytical approaches, contrasting with our use of computerised tests.

The implementation of computerised cognitive batteries, specifically the CANTAB, offers enhanced measurement precision and consistency, reducing examiner-related variability and potentially increasing the sensitivity to detect subtle cognitive changes. The CANTAB-PAL has demonstrated effectiveness in episodic memory assessment, having established robust psychometric properties across diagnostic, cultural and linguistic populations (Barnett et al., 2016). The test exhibits strong temporal reliability (Barnett et al., 2016) and has demonstrated strong correlations with both conventional neuropsychological tests (Matos Gonçalves et al., 2018) and functional assessment metrics (Barnett et al., 2016). Additionally, the CANTAB-DMS paradigm has been validated in clinical populations, including individuals with mild cognitive impairment, dementias and Parkinson’s disease (Cabral Soares et al., 2014; Campos-Magdaleno et al., 2021; Fowler et al., 1995; Sahakian et al., 1988), supporting its use in cognitive assessment. The enhanced psychometric properties inherent in these computerised assessments may account for the detection of significant interaction effects between the aperiodic exponent and age on episodic and visual short-term memory in our investigation, contrasting with the findings reported in studies utilising traditional verbal and pen-and-paper assessment methods.

Interestingly, the interaction effect between age and the aperiodic exponent was not a significant predictor in the spatial working memory task. Research has shown that performance on the PAL and DMS demonstrates selective sensitivity to medial temporal lobe (MTL) functionality (Owen et al., 1995; Torgersen et al., 2010), particularly hippocampal integrity, which supports both episodic memory processes (Dickerson & Eichenbaum, 2010; Nyberg et al., 2000) and short-term retention of visual information (Barense et al., 2007; Lee et al., 2012; Olson et al., 2006; Warren et al., 2012). In contrast, the spatial working memory task predominantly relies on distributed frontoparietal networks (Courtney et al., 1998; Curtis, 2006; D’Esposito et al., 1998; Schweinsburg et al., 2005; van Asselen et al., 2006). This raises the possibility that the exponent may specifically index the vulnerability or resilience of MTL-dependent cognitive processes to age-related decline, rather than serving as a general marker of cognitive ageing across all domains, though further investigation utilising multimodal neuroimaging approaches is warranted to understand the precise neurophysiological mechanisms underlying these domain-specific effects.

Additionally, the observed interaction effects between the exponent and age demonstrated topographically distinct patterns across regions, implicating the fronto-central electrode cluster for visual short-term memory accuracy and the parieto-occipital electrode cluster for visuospatial episodic memory errors. A prior study by Pathania et al. (2022) found that the exponent within frontal regions served as a significant mediating factor in age-related attentional performance differences between younger and older adults, although statistical comparisons across brain regions were not conducted.

Similarly, while the present study may suggest that distinct neural networks contribute to the moderating role of the aperiodic exponent in age-related changes in memory processes, our study design and analyses did not directly compare electrode clusters. Future investigations employing statistical comparisons across scalp regions are warranted to investigate the spatial distribution of age-exponent interactions and their differential impact on distinct memory processes.

Furthermore, HLR analyses revealed no significant direct associations between the aperiodic exponent and memory performance. This finding contrasts with recent evidence supporting the predictive value of the exponent in cognitive performance, particularly in older adult populations (Smith et al., 2023) and across the lifespan (Finley et al., 2024). However, other investigations have similarly failed to demonstrate such associations, suggesting complex underlying relationships (Cesnaite et al., 2023; Tröndle et al., 2023). The current findings indicate that the exponent’s influence on cognitive performance may be contextually dependent, manifesting primarily through interactions with age-related factors rather than through direct effects, though further research is required to clarify these findings.

The present study also revealed no significant direct associations between IAF and memory performance across tasks or a significant moderation effect of IAF and age on memory performance. This finding contrasts with recent evidence demonstrating IAF’s predictive value for cognitive performance, such as sustained attention in older adults, as per our previous study (Campbell et al., 2024), and longitudinal cognitive decline across middle to late adulthood (Finley et al., 2024).

Additionally, prior research has established positive associations between IAF and interference resolution in working memory performance within older adults (Cesnaite et al., 2023), making the absence of significant relationships in our study particularly noteworthy. Again, this may be due to methodological differences. The substantially larger sample size employed in the aforementioned study may have provided enhanced statistical power for detecting subtle associations with IAF within this age demographic. Furthermore, methodological differences in working memory paradigms warrant consideration, as Cesnaite et al. (2023) investigated interference resolution in working memory, whereas the current study examined errors in spatial working memory among other memory systems. The spatial distribution of observed effects also differed, with the prior study identifying primarily temporal lobe associations, while the present study focused exclusively on fronto-central and parieto-occipital electrode clusters. Future investigations should address these discrepancies by incorporating multiple memory paradigms, expanded spatial sampling, and larger cohort sizes.

### 4.1 Limitations

There are several limitations inherent within our study that require discussion. Firstly, the cross-sectional nature of our study design precluded the determination of causal relationships as well as any longitudinal relationships between neurophysiological parameters and cognitive performance. The sample demographics demonstrate a substantial gender bias with female predominance, restricting generalisability across gender groups. Our cohort’s homogeneity, comprising highly functional and well-educated older adults, may also not adequately represent the broader ageing population, particularly individuals experiencing cognitive decline. Additional methodological limitations include the restricted spatial sampling, with the present study focusing exclusively on fronto-central and parieto-occipital electrode clusters. Future research would benefit from comprehensive coverage of other cortical areas or statistical comparisons between clusters, as well as additional frequency bands. Additionally, the absence of task-based EEG restricts analysis of state-dependent variations in neural dynamics. Finally, the single-modality neuroimaging approach constrains integration with other age-related neurobiological alterations documented in the literature.

## 5.0 Conclusion

The present study uncovered key relationships among rsEEG aperiodic exponents, age, and memory function in older adults aged 50-84 years. The findings demonstrate a significant moderating effect of the exponent on age-related performance in both episodic and visual short-term memory systems.

Notably, for individuals with higher exponent values, age was not significantly associated with episodic or visual short-term memory performance, whereas those with average and lower exponent values showed poorer episodic and visual short-term memory performance with advancing age. This pattern suggests that lower and average aperiodic exponents may reflect a marker of decrement in age-related memory performance, whereas steeper exponents, potentially reflecting an optimised E/I balance within neural circuits, may index a protective mechanism against age-related memory decline. The present findings provide new insights into the neurophysiological mechanisms underlying cognitive ageing and identify the aperiodic exponent as a potential biomarker of individual differences in age-related memory performance trajectories. Finally, our study establishes a theoretical framework for investigating neural mechanisms that may offer protection against age-related cognitive decline, potentially informing the development of interventional approaches targeting preserved cognitive function in older populations and advances the understanding of the complex interplay between neurophysiological parameters and cognitive ageing.

## CRediT Authorship Contribution Statement

**Alicia J. Campbell:** Conceptualisation, Methodology, Software, Formal Analysis, Investigation, Data Curation, Writing-Original Draft, Visualisation; **Toomas Erik Anijärv:** Software, Data Curation, Writing – Review & Editing; **Mikael Johansson:** Conceptualisation, Methodology, Writing – Review & Editing; **Thomas Pace:** Writing – Review & Editing**; Jim Lagopoulos:** Writing – Review & Editing; **Daniel F. Hermens:** Writing – Review & Editing, Supervision; **Jacob M. Levenstein:** Writing – Review & Editing, Supervision; **Sophie C. Andrews:** Conceptualisation, Methodology, Formal Analysis, Writing – Review & Editing, Supervision.

## Data Availability

The scripts for EEG preprocessing and spectral analysis are publicly available at https://github.com/aliciajcampbell/EEG-notebooks, while the scripts for statistical analyses are publicly available at https://github.com/aliciajcampbell/LEISURE-EEG-cog-analyses.

