## Supplementary Materials for "Resting-State EEG Aperiodic Exponent Moderates the Association Between Age and Memory Performance in Older Adults"

**Supplementary Material.**

**Table S1:** Skewness and kurtosis values for all variables, including raw and outlier corrected values.

| | Variable | Skewness | Kurtosis |
|---|---|---|---|
| **Raw values** | Age | 0.032 | -0.991 |
| | Gender | -1.765 | 1.141 |
| | Education | 2.772 | 17.854 |
| | Handedness | -2.137 | 2.627 |
| | DMS PCAD | -0.644 | -0.092 |
| | PAL TEA | 1.014 | 0.043 |
| | SWM BE | -0.401 | 0.639 |
| | Fronto-central IAF | 0.483 | 0.619 |
| | Fronto-central exponent | 0.052 | 0.440 |
| | Parieto-occipital IAF | -0.097 | -0.449 |
| | Parieto-occipital exponent | -0.124 | -0.328 |
| **Corrected values** | Education | -0.425 | -1.013 |
| | SWM BE | 0.176 | -0.521 |
| | Fronto-central IAF | -0.747 | -0.182 |

DMS PCAD, Delayed Match to Sample percent correct (all delays, visual short-term memory); PAL TEA, Paired Associates Learning total errors (adjusted, visuospatial episodic memory); SWM BE, Spatial Working Memory between errors; IAF, Individual alpha peak frequency; variables corrected for outliers using z-score standard deviation transformation method (Corrected values).

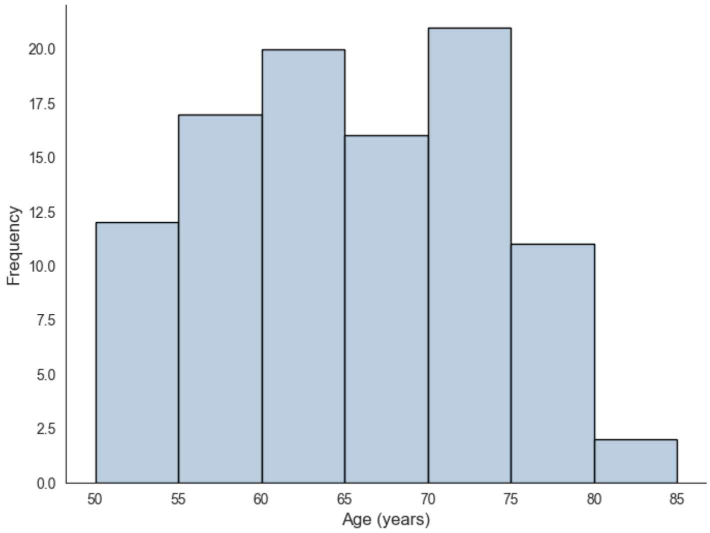


**Figure S1.** Histogram of participant ages (N = 99), binned in 5-year intervals from 50 to 85 years.

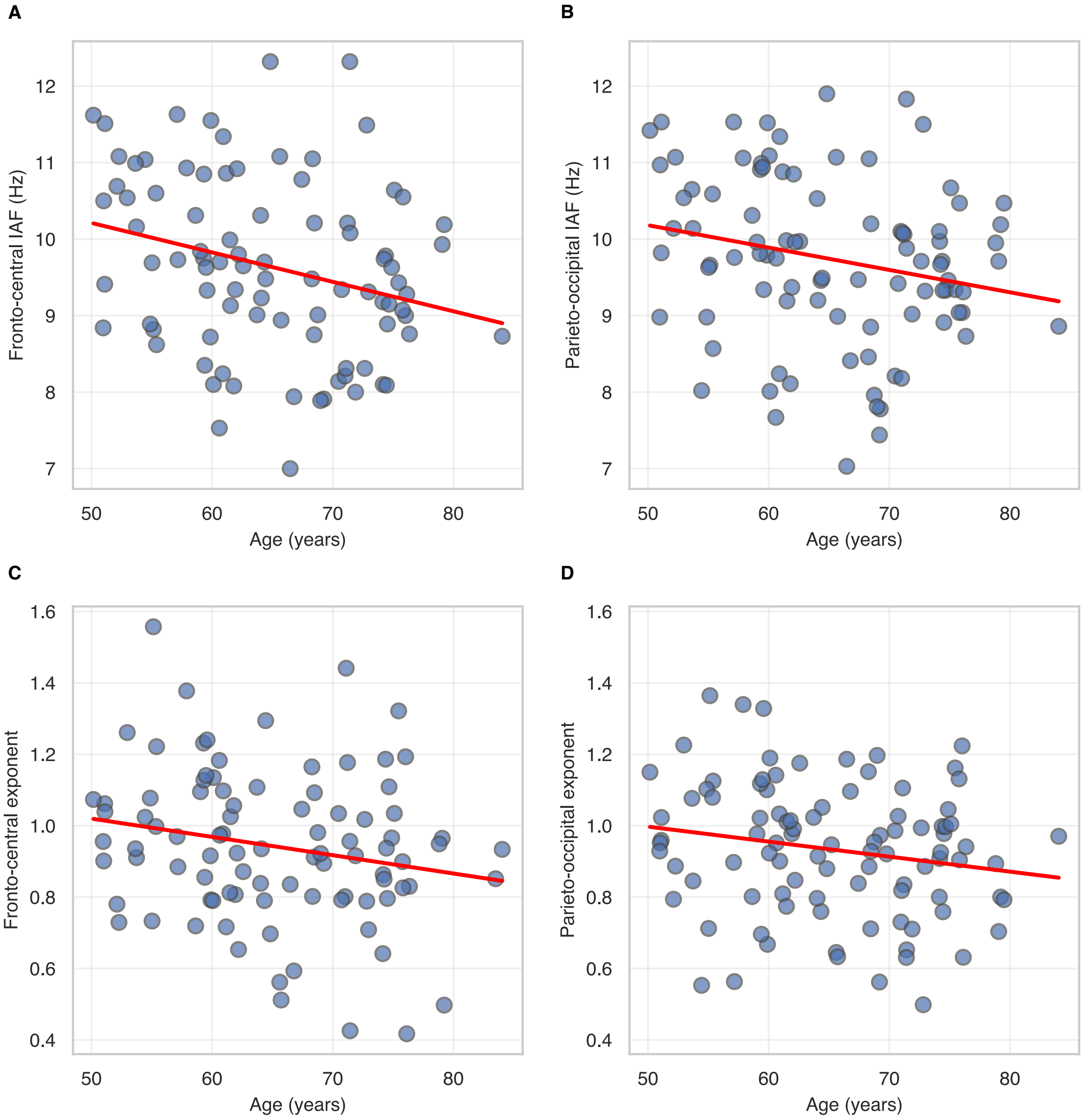


**Figure S2.** Scatterplots showing the association between age and resting-state EEG markers. Panels (A) and (B) depict IAF at fronto-central and parieto-occipital electrodes, respectively. Panels (C) and (D) show the aperiodic exponent at fronto-central and parieto-occipital sites. Red lines represent linear regression fits. IAF, Individual alpha peak frequency.

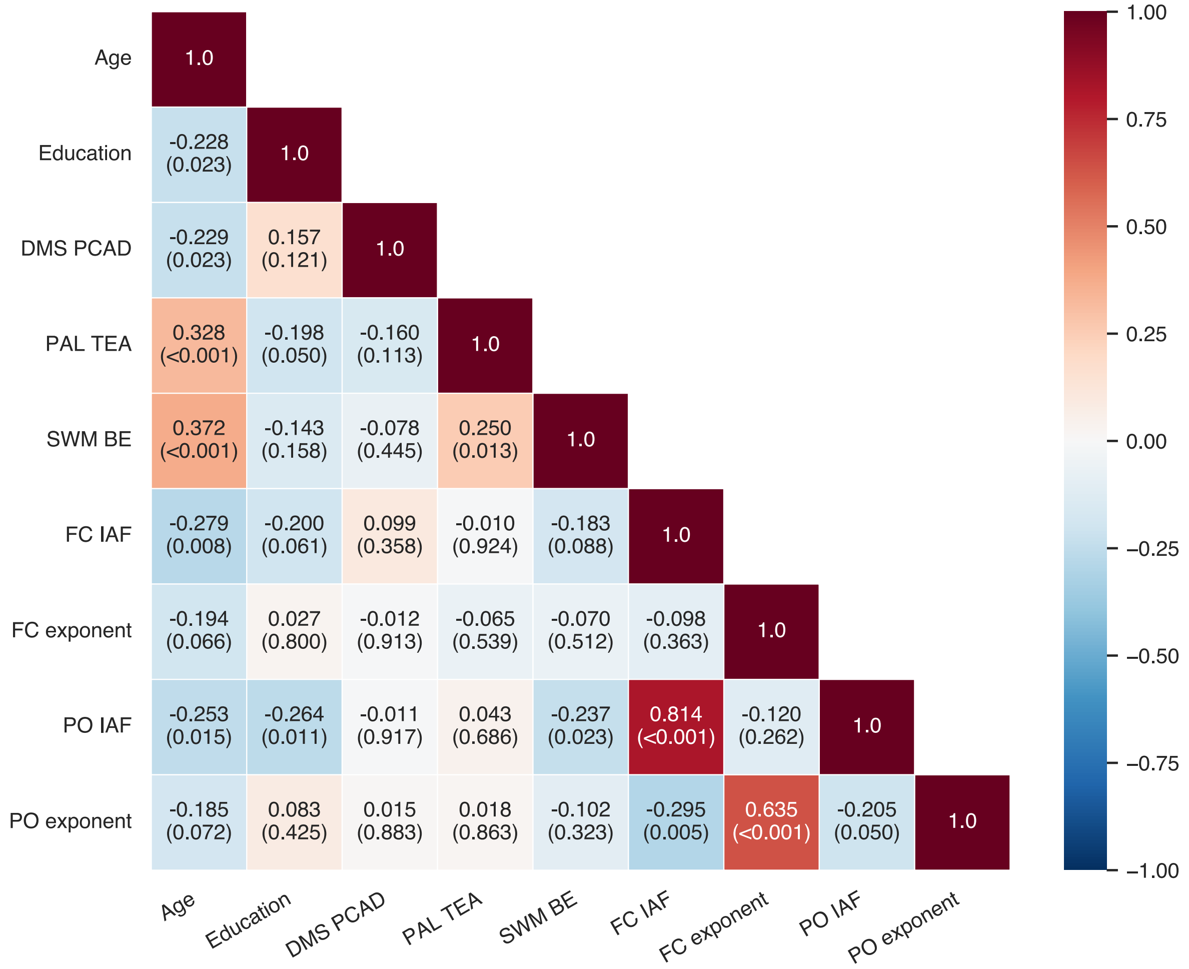


**Figure S3.** Spearman rho correlation coefficients (ρ) (coloured by magnitude) between performance across memory tasks, age, education and resting-state EEG measures, with p-values provided in parentheses; p-values represent uncorrected values. DMS PCAD, Delayed Match to Sample percent correct (all delays, visual short-term memory); PAL TEA, Paired Associates Learning total errors (adjusted, visuospatial episodic memory); SWM BE, Spatial Working Memory between errors; FC, fronto-central; PO, parieto-occipital; IAF, Individual alpha peak frequency.

**Table S2.** Hierarchical linear regression models predicting memory task performance from main effects of fronto-central IAF and aperiodic exponent values and interactions with age, including demographic covariates as step 1 regressors.

| Dependent Variable | Predictors | Step | β | *p*(β) | $R^2$ | F | *p*(F) | $ΔR^2$ | ΔF | *p*(ΔF) |
|---|---|---|---|---|---|---|---|---|---|---|
| DMS PCAD | Age | 1 | -.236 | .032* | .097 | 2.235 | .072 | | | |
| | Gender | | .014 | .895 | | | | | | |
| | Education | | .079 | .467 | | | | | | |
| | Handedness | | -.149 | .172 | | | | | | |
| | IAF | 2 | .036 | .756 | .098 | 1.788 | .124 | .001 | .097 | .756 |
| | Age × IAF | 3 | -.126 | .248 | .113 | 1.722 | .126 | .015 | 1.352 | .248 |
| DMS PCAD | Age | 1 | -.220 | .044* | .084 | 1.981 | .105 | | | |
| | Gender | | -.029 | .788 | | | | | | |
| | Education | | .057 | .597 | | | | | | |
| | Handedness | | -.138 | .203 | | | | | | |
| | Exponent | 2 | -.099 | .350 | .094 | 1.759 | .130 | .009 | .881 | .350 |
| | Age × exponent | 3 | .221 | .035 | .141 | 2.294 | .042* | .047 | 4.597 | .035* |

| | | | | | | | | | | |
|---|---|---|---|---|---|---|---|---|---|---|
| PAL TEA | Age | 1 | .280 | .011* | .118 | 2.776 | .032* | | | |
| | Gender | | -.029 | .789 | | | | | | |
| | Education | | -.111 | .305 | | | | | | |
| | Handedness | | .096 | .374 | | | | | | |
| | IAF | 2 | .019 | .872 | .118 | 2.200 | .062 | <.001 | .026 | .872 |
| | Age × IAF | 3 | .074 | .497 | .123 | 1.899 | .091 | .005 | .466 | .497 |
| PAL TEA | Age | 1 | .291 | .007** | .124 | 3.036 | .022* | | | |
| | Gender | | -.023 | .824 | | | | | | |
| | Education | | -.102 | .336 | | | | | | |
| | Handedness | | .092 | .382 | | | | | | |
| | Exponent | 2 | -.034 | .745 | .125 | 2.425 | .042* | .001 | .107 | .745 |
| | Age × exponent | 3 | -.088 | .398 | .132 | 2.134 | .058 | .007 | .721 | .398 |
| SWM BE | Age | 1 | .445 | <.001*** | .207 | 5.403 | .001** | | | |
| | Gender | | -.101 | .323 | | | | | | |
| | Education | | -.014 | .891 | | | | | | |
| | Handedness | | -.005 | .957 | | | | | | |
| | IAF | 2 | -.061 | .581 | .210 | 4.348 | .001** | .003 | .306 | .581 |
| | Age × IAF | 3 | .050 | .629 | .212 | 3.628 | .003** | .002 | .235 | .629 |
| SWM BE | Age | 1 | .443 | <.001*** | .204 | 5.494 | .001** | | | |
| | Gender | | -.077 | .443 | | | | | | |
| | Education | | -.006 | .951 | | | | | | |
| | Handedness | | -.011 | .911 | | | | | | |
| | Exponent | 2 | .025 | .802 | .204 | 4.36 | .001** | .001 | .063 | .802 |
| | Age × exponent | 3 | -.030 | .703 | .205 | 3.621 | .003** | .001 | .147 | .703 |

β, standardised beta; "Step" outlines which level of the HLR analyses the variables were entered; DMS PCAD, Delayed Match to Sample percent correct (all delays, visual short-term memory); PAL TEA, Paired Associates Learning total errors (adjusted, visuospatial episodic memory); SWM BE, Spatial Working Memory between errors; IAF, Individual alpha peak frequency. *p<.05, **p<.01, ***p<.001.

**Table S3**. Hierarchical linear regression models predicting memory task performance from main effects of parieto-occipital IAF and aperiodic exponent values and interactions with age, including demographic covariates as step 1 regressors.

| Dependent Variable | Predictors | Step | β | *p*(β) | $R^2$ | F | *p*(F) | $\Delta R^2$ | ΔF | *p*(ΔF) |
|---|---|---|---|---|---|---|---|---|---|---|
| DMS PCAD | Age | 1 | -.254 | .017* | .109 | 2.663 | .038* | | | |
| | Gender | | -.001 | .994 | | | | | | |
| | Education | | .079 | .453 | | | | | | |
| | Handedness | | -.151 | .153 | | | | | | |
| | IAF | 2 | -.035 | .758 | .110 | 2.128 | .070 | .001 | .096 | .758 |
| | Age × IAF | 3 | -.053 | .617 | .113 | 1.800 | .109 | .003 | .252 | .617 |
| DMS PCAD | Age | 1 | -.250 | .017* | .107 | 2.686 | .036* | | | |
| | Gender | | -.023 | .824 | | | | | | |
| | Education | | .085 | .410 | | | | | | |

| | | | | | | | | | | |
|---|---|---|---|---|---|---|---|---|---|---|
| | Handedness | | -.141 | .174 | | | | | | |
| | Exponent | 2 | -.056 | .584 | .110 | 2.192 | .062 | .003 | .301 | .584 |
| | Age × exponent | 3 | .183 | .075 | .141 | 2.415 | .033* | .032 | 3.254 | .075 |
| PAL TEA | Age | 1 | .320 | .002** | .143 | 3.620 | .009** | | | |
| | Gender | | -.074 | .474 | | | | | | |
| | Education | | -.093 | .368 | | | | | | |
| | Handedness | | .110 | .287 | | | | | | |
| | IAF | 2 | .113 | .313 | .153 | 3.103 | .013* | .010 | 1.032 | .313 |
| | Age × IAF | 3 | .115 | .269 | .165 | 2.800 | .016* | .012 | 1.238 | .269 |
| PAL TEA | Age | 1 | .314 | .002** | .141 | 3.692 | .008** | | | |
| | Gender | | -.072 | .479 | | | | | | |
| | Education | | -.106 | .296 | | | | | | |
| | Handedness | | .103 | .309 | | | | | | |
| | Exponent | 2 | <.001 | .997 | .141 | 2.921 | .017* | <.001 | <.001 | .997 |
| | Age × exponent | 3 | -.220 | .028 | .187 | 3.377 | .005** | .046 | 5.000 | .028* |
| SWM BE | Age | 1 | .446 | <.001*** | .208 | 5.706 | <.001*** | | | |
| | Gender | | -.072 | .469 | | | | | | |
| | Education | | -.022 | .825 | | | | | | |
| | Handedness | | -.006 | .953 | | | | | | |
| | IAF | 2 | -.143 | .180 | .224 | 4.973 | <.001*** | .016 | 1.825 | .180 |
| | Age × IAF | 3 | .166 | .094 | .250 | 4.712 | <.001*** | .025 | 2.869 | .094 |
| SWM BE | Age | 1 | .432 | <.001*** | .205 | 5.818 | <.001*** | | | |
| | Gender | | -.111 | .256 | | | | | | |
| | Education | | -.039 | .686 | | | | | | |
| | Handedness | | -.001 | .990 | | | | | | |
| | Exponent | 2 | .009 | .924 | .206 | 4.605 | .001** | <.001 | .009 | .924 |
| | Age × exponent | 3 | -.013 | .891 | .206 | 3.798 | .002** | <.001 | .019 | .891 |

β, standardised beta; "Step" outlines which level of the HLR analyses the variables were entered; DMS PCAD, Delayed Match to Sample percent correct (all delays, visual short-term memory); PAL TEA, Paired Associates Learning total errors (adjusted, visuospatial episodic memory); SWM BE, Spatial Working Memory between errors; IAF, Individual alpha peak frequency. **p*<.05, ***p*<.01, ****p*<.001.

**Aperiodic offset analyses**

Additional analyses were conducted on the aperiodic offset parameter extracted using the specparam toolbox. The offset was estimated from the same parameterisation used to derive the aperiodic exponent (fixed aperiodic mode; 1–30 Hz fitting range) and averaged across fronto-central and parieto-occipital electrode clusters. Moreover, exclusions of offset data were identical to exponent data.

Statistical procedures were identical to those described in the main manuscript. Bivariate correlations were analysed between the aperiodic offset and age, education, other rsEEG measures (IAF and exponent at fronto-central and parieto-occipital electrodes), and memory task performance (DMS PCAD, PAL TEA, SWM BE) using Spearman's rank correlation (two-tailed). To control for multiple comparisons, p-values were corrected using the FDR method at $q$ = .05 for a total of 18 pairwise comparisons.

Neither fronto-central ($N$=91, M=.49, SD=.25, min=-.03, max=1.19) nor parieto-occipital ($N$=95, M=.47, SD=.26, min=-.10, max=1.25) offset was significantly associated with age, education, or memory performance after FDR correction (Table S4).

However, both offset measures showed positive associations with their corresponding regional aperiodic exponents cross-regional associations with exponents in the opposite electrode cluster. No significant correlations were observed between offset and IAF in either region.

**Table S4.** Bivariate Spearman correlations (ρ) between fronto-central and parieto-occipital aperiodic offset, age, education, rsEEG measures, and memory task performance.

| **Variable 1** | **Variable 2** | **ρ** | ***p*** | ***p* (adjusted)** |
|---|---|---|---|---|
| **Fronto-central offset** | Age | -.211 | .044 | .133 |
| **Fronto-central offset** | Education | .030 | .774 | .946 |
| **Fronto-central offset** | Fronto-central IAF | -.007 | .946 | .946 |
| **Fronto-central offset** | Parieto-occipital IAF | .014 | .899 | .946 |
| **Fronto-central offset** | Fronto-central exponent | .533 | <.001*** | <.001*** |
| **Fronto-central offset** | Parieto-occipital exponent | .440 | <.001*** | <.001*** |
| **Fronto-central offset** | DMS PCAD | -.078 | .462 | .940 |
| **Fronto-central offset** | PAL TEA | -.012 | .908 | .946 |
| **Fronto-central offset** | SWM BE | -.052 | .627 | .940 |
| **Parieto-occipital offset** | Age | -.220 | .032* | .117 |
| **Parieto-occipital offset** | Education | .071 | .495 | .940 |
| **Parieto-occipital offset** | Fronto-central IAF | -.059 | .586 | .940 |
| **Parieto-occipital offset** | Parieto-occipital IAF | -.040 | .707 | .946 |
| **Parieto-occipital offset** | Fronto-central exponent | .379 | <.001*** | <.001*** |
| **Parieto-occipital offset** | Parieto-occipital exponent | .546 | <.001*** | <.001*** |
| **Parieto-occipital offset** | DMS PCAD | -.010 | .926 | .946 |
| **Parieto-occipital offset** | PAL TEA | .059 | .568 | .940 |
| **Parieto-occipital offset** | SWM BE | -.051 | .625 | .940 |

*p* (adjusted), adjusted using false discovery rate; ρ, spearman rho correlation coefficients; DMS PCAD, Delayed Match to Sample percent correct (all delays, visual short-term memory); PAL TEA, Paired Associates Learning total errors (adjusted, visuospatial episodic memory); SWM BE, Spatial Working Memory between errors; IAF, Individual alpha peak frequency. **p*<.05, ***p*<.01, ****p*<.001.

Additionally, HLR models were performed to examine associations between offset and memory task performance. Each HLR model included age, gender, education, and handedness as covariates in Step 1, the offset in Step 2, and the Age × Offset interaction term in Step 3. Improvements in model fit were evaluated using the F-change statistic (ΔF).

After controlling for age, gender, education, and handedness, neither fronto-central nor parieto-occipital offset was significantly associated with performance on any of the memory tasks (Table S5). Additionally, no significant interaction effects between age and offset were observed for either region.

**Table S5**. Hierarchical linear regression models predicting memory task performance from main effects of fronto-central and parieto-occipital aperiodic offset values and interactions with age, including demographic covariates as step 1 regressors.

| **Dependent Variable** | **Predictors** | **Step** | **β** | ***p*(β)** | **$R^2$** | **F** | ***p*(F)** | **$\Delta R^2$** | **ΔF** | ***p*(ΔF)** |
|---|---|---|---|---|---|---|---|---|---|---|
| DMS PCAD | Age | 1 | -.220 | .044* | .084 | 1.981 | .105 | | | |
| | Gender | | -.029 | .788 | | | | | | |
| | Education | | .057 | .597 | | | | | | |
| | Handedness | | -.138 | .203 | | | | | | |
| | Offset | 2 | -.140 | .191 | .103 | 1.946 | .095 | .018 | 1.736 | .191 |
| | Age × fronto-central offset | 3 | -.137 | .836 | .103 | 1.610 | .154 | <.001 | .043 | .836 |

| | | | | | | | | | | |
|---|---|---|---|---|---|---|---|---|---|---|
| DMS PCAD | Age | 1 | -.250 | .017* | .107 | 2.686 | .036* | | | |
| | Gender | | -.023 | .824 | | | | | | |
| | Education | | .085 | .410 | | | | | | |
| | Handedness | | -.141 | .174 | | | | | | |
| | Offset | 2 | -.108 | .297 | .118 | 2.371 | .045* | .011 | 1.100 | .297 |
| | Age × parieto-occipital offset | 3 | -.107 | .920 | .118 | 1.956 | .081 | <.001 | .010 | .920 |
| PAL TEA | Age | 1 | .291 | .007** | .124 | 3.036 | .022* | | | |
| | Gender | | -.023 | .824 | | | | | | |
| | Education | | .102 | .336 | | | | | | |
| | Handedness | | .092 | .382 | | | | | | |
| | Offset | 2 | .065 | .538 | .128 | 2.488 | .037* | .004 | .383 | .538 |
| | Age × fronto-central offset | 3 | <-.001 | .998 | .128 | 2.049 | .068 | <.001 | <.001 | .998 |
| PAL TEA | Age | 1 | .314 | .002** | .141 | 3.692 | .008** | | | |
| | Gender | | -.072 | .479 | | | | | | |
| | Education | | -.106 | .296 | | | | | | |
| | Handedness | | .103 | .309 | | | | | | |
| | Offset | 2 | .108 | .291 | .152 | 3.184 | .011* | .011 | 1.130 | .291 |
| | Age × parieto-occipital offset | 3 | -.144 | .149 | .172 | 3.039 | .010* | .020 | 2.118 | .149 |
| SWM BE | Age | 1 | .443 | <.001*** | .204 | 5.494 | .001** | | | |
| | Gender | | -.077 | .443 | | | | | | |
| | Education | | -.006 | .951 | | | | | | |
| | Handedness | | -.011 | .911 | | | | | | |
| | Offset | 2 | .072 | .474 | .208 | 4.474 | .001** | .005 | .518 | .474 |
| | Age × fronto-central offset | 3 | -.063 | .521 | .212 | 3.772 | .002** | .004 | .416 | .521 |
| SWM BE | Age | 1 | .432 | <.001*** | .205 | 5.818 | <.001*** | | | |
| | Gender | | -.111 | .256 | | | | | | |
| | Education | | -.039 | .686 | | | | | | |
| | Handedness | | -.001 | .990 | | | | | | |
| | Offset | 2 | .082 | .401 | .212 | 4.782 | .001** | .006 | .712 | .401 |
| | Age × parieto-occipital offset | 3 | -.103 | .285 | .222 | 4.184 | .001** | .010 | 1.156 | .285 |

β, standardised beta; "Step" outlines which level of the HLR analyses the variables were entered; DMS PCAD, Delayed Match to Sample percent correct (all delays, visual short-term memory); PAL TEA, Paired Associates Learning total errors (adjusted, visuospatial episodic memory); SWM BE, Spatial Working Memory between errors. *$p$<.05, **$p$<.01, ***$p$<.001.
